# Effects of High-Grain Diet with Buffering Agent on the Milk Protein Synthesis in Lactating Goats

**DOI:** 10.1101/2020.05.12.091173

**Authors:** Meilin He, Xintian Nie, Huanhuan Wang, Shuping Yan, Yuanshu Zhang

**Author notes:** To whom correspondence should be addressed: Yuanshu Zhang: The Key Laboratory of Animal Physiology and Biochemistry, Ministry of Agriculture, Nanjing Agricultural University, Nanjing 210095, People’s Republic of China.

## Abstract

Feeding of straw as main roughage with numerous high-grain diets improves the performance of ruminants but it can easily lead to subacute ruminal acidosis. In recent years, buffering agent is applied to prevent the acid poisoning of ruminants and improve the production performance of ruminants in animal husbandry. it is necessary to understand feeding high-grain diet with buffering agent which transport carriers amino acids mainly take amino acids into the mammary gland and the signal mechanism of amino acids in the mammary gland synthesize milk proteins. To gain insight on the effects of a high-grain diet with buffering agent on the amino acids in the jugular blood, and the effects of amino acids on the synthesis of milk protein, commercial kit and high performance liquid chromatography (HPLC) were applied to determine the concentration of amino acids of jugular blood samples, quantitative real-time PCR, comparative proteomic approach and western blot were employed to investigate proteins differentially expressed in mammary tissues and the mechanism of amino acids on the synthesis of milk protein in mammary gland of lactating dairy goats fed high-grain diet with buffering agent or only high-grain diet.

Results showed that feeding high-grain diet with buffering agent to lactating dairy goats could outstanding increase amino acid content of jugular blood (p<0.05), and mRNA transcriptional level of amino acid transporters in the mammary gland were also increased; the CSN2 and LF protein expression level were significant higher by 2-DE technique, MALDI-TOF/TOF proteomics analyzer and western blot analysis further validated in mammary of lactating dairy goats compared with high-grain group; the research on the mechanism of milk protein synthesis increasing suggested that it was related to the activation of mTOR pathway signaling.

Feeding of high-grain diet with buffering agent promoted the jugular vein blood of amino acids concentration, and more amino acids flowed into the mammary. In addition, milk protein synthesis was increased and the increase of milk protein synthesis was related to the activation of mTOR pathway signalling.

## Background

Due to reduction of per capita arable land, the degradation of grasslands, and shortage of green fodder resources and poor quality, the current feeding practices in the dairy industry improving feed quality by feeding high-grain diets. However, many studies have confirmed that feeding of straw as main roughage with numerous high-grain diets improves the performance of ruminants but it can easily lead to subacute ruminal acidosis (SARA) (1,2). SARA has observed a decline in ruminal pH below 5.6 fed such ration for 3-5 h (3).

Buffering agent is a chemical that enhances the acid-base buffer capacity of a solution. In recent years, it is applied to prevent the acid poisoning of ruminants and improve the production performance of ruminants in animal husbandry. A comprehensive study of three American universities has shown that adding 1.5% sodium bicarbonate and 0.8% magnesium oxide in the diets of early and mid-term lactating dairy cows can cause milk production and milk fat levels to increase significantly (4).

Kurokawa et al. added 80g of sodium bicarbonate every day when feeding the lactating dairy cows with salt which was less than 20% of standard, compared with the control group, the milk yield was 5.1% and the butterfat percentage was 0.15% below the standard, and compared with the control group, the standard milk was the increased by 5.1% and the rate of butterfat increased by 0.15% (5). In order to maintain ruminal pH in lactating dairy goats, buffering agent was added to the high-grain diets.

Milk protein is one of the determinants of milk quality, and milk proteins in the lactating ruminant mammary gland are primarily synthesized from circulating plasma amino acids (6). Previous studies have shown that amino acids can be taken into the mammary gland by the mammary epithelial cells through the transport carriers (7–9). Therefore, it is necessary to understand feeding high-grain diet with buffering agent which transport carriers amino acids mainly take amino acids into the mammary gland and the signal mechanism of amino acids in the mammary gland synthesize milk proteins. In this way, it can alleviate SARA’s symptoms and improve the milk protein production and lay a foundation for future research.

## Methods

### Experimental animals

A total of twelve healthy multiparous mid-lactating goats (body weight, 38 ±8 kg, mean ± SEM, 3-5 weeks post-partum) at the age of 2-3 years were used in experiments. They were housed in individual stalls in a standard animal feeding house at Nanjing Agricultural University (Nanjing, China). Goats were randomly divided into two groups: high-grain group (HG, concentrate: forage = 60:40) and buffering agent group (BG, concentrate: forage = 60:40 with 10g C_4_H_7_NaO_2_ and 10g NaHCO_3_), six in each group. Dietary C_4_H_7_NaO_2_ and NaHCO_3_ were obtained from Nanjing Jian cheng Bioengineering Institute, China. The ingredients and nutritional composition of the diets are presented in Table 1. The goats were fitted with a rumen fistula and hepatic catheters two weeks before the experiment and were ensured that they recovered from the surgery. Animals were monitored for 2 weeks after surgery. Sterilized heparin saline (500 IU/ml, 0.3 ml/time) was administered at 8-hour intervals every day until the end of the experiment to prevent catheters from becoming blocked. During the experimental period of 20 weeks, goats were fed two times daily at 8.00 and 18.00, had free access to fresh water, and the feed amount met or exceeded the animal’s nutritional requirements. The Institutional Animal Care and Use Committee of Nanjing Agricultural University (Nanjing, People’s Republic of China) approved all of the procedures (surgical procedures and care of goats).

**Table 1.** Ingredients and nutritional composition of the diets.

| Concentrate: Forage ratio 60:40 |  |  |  |
| --- | --- | --- | --- |
| Ingredient (%) |  | Nutrient levels<sup>b</sup> |  |
| Leymus chinensis | 27.00 | Net energy/(MJ.kg<sup>-1</sup>) | 6.71 |
| Alfalfa silage | 13.00 | Crude protein/% | 16.92 |
| Corn | 23.24 | Neutral detergent fiber/% | 31.45 |
| Wheat bran | 20.77 | Acid detergent fiber/% | 17.56 |
| Soybean meal | 13.67 | Calcium/% | 0.89 |
| Limestone | 1.42 | Phosphorus/% | 0.46 |
| NaCl | 0.30 |  |  |
| Premix<sup>a</sup> | 0.60 |  |  |
| Total | 100.00 |  |  |

a. Provided per kg of diet: VA 6000IU/kg, VD 2500IU/kg, VE 80mg/kg, Cu 6.25 mg/kg, Fe 62.5 mg/kg, Zn 62.5 mg/kg, Mn 50mg/kg, I 0.125 mg/kg, Co 0.125 mg/kg.
b. Nutrient levels were according to National Research Council (NRC,2001).

### Analysis of total amino acids

At the 20^th^ week, blood samples were collected from the jugular blood in 10 mL vacuum tubes containing sodium heparin. Blood was centrifuged at 3,000 × g for 15 min to separate plasma. The total amino acids concentration were determined by a Total Amino Acid assay kit (catalog no. A026, Jiancheng, Nanjing, China). The procedures were performed according to the manufacturer’s instructions.

### Analyses of amino acids profile by HPLC

Free amino acids of jugular blood samples were determined by high performance liquid chromatography (HPLC), it was performed as previously described by Shen et al (10). The HPLC system consisted of: Agilent1100 high-performance liquid chromatograph system (Agilent Technologies, Waldbronn, Germany); scanning fluorescence detector (excitation 340 nm, emission 450 nm); chromatographic column (XTerra^®^MS C18, 4.6mm × 250mm, 5μm), which was purchased from waters (Waters Co., Milford, MA, USA). 20 kinds of standard amino acids (Aldrich chemical company) were given by professor Holey, and the purity of these amino acids were greater than 98%. The three-dimensional flow phase (solution A, methanol; solution B, acetonitrile; solution C, 10mmol/L phosphate buffer containing 0.3% tetrahydrofuran) was adopted. The gradient program was referred to Table 2, and oven temperature was 40 °C, the injection volume was 20 μL. The plasma samples were mixing with acetonitrile by 1:2(v/v), and were placed at 4°C for 30 min, then they were centrifuged at 12000 rpm for 30 min, and the supernatant fluids were collected for AA analysis. High pressure liquid chromatography analysis was performed after automatic pre-column derivatization with O-phthaldialdehyde (OPA) (11).

**Table 2.** Gradient elution program of RP-HPLC.

| Time (min) | Solution A (%) | Solution B (%) | Solution C (%) | Flow rate (mL/min) |
| --- | --- | --- | --- | --- |
| 0 | 85 | 6 | 9 | 1 |
| 10 | 80 | 8 | 12 | 1 |
| 25 | 70 | 15 | 15 | 1 |
| 40 | 45 | 25 | 30 | 1 |
| 50 | 45 | 25 | 30 | 1 |

### Quantitative Real-Time PCR (qRT-PCR)

After 20 weeks, goats were slaughtered after overnight fasting. All goats were killed with neck vein injections of xylazine (0.5 mg (kg body weight)^−1^; Xylosol; Ogris Pharme, Wels, Austria) and pentobarbital (50 mg (kg body weight)^−1^; Release; WDT, Garbsen, Germany). After slaughter, mammary tissue was collected and washed twice with cold physiological saline (0.9% NaCl) to remove blood and other contaminants. Mammary tissue samples were used for RNA and protein extraction. Total RNA was extracted from each mammary tissue sample using the TRIzol reagent (Invitrogen, USA) according to the manufacturer’s specifications and then reverse-transcribed into cDNA using commercial kits (Vazyme, Nanjing, China). All PCR primers were synthesized by Generay Company (Shanghai, China), and the primer sequences are listed in Table 3. PCR was performed using the AceQ qPCR SYBR Green Master Mix kit (Vazyme, Nanjing, China) and the MyiQ2 Real-time PCR system (Bio-Rad, USA) with the following cycling conditions: 95°C for 2 min, 40 cycles of 95°C for 15 sec and 60°C for 30 sec. Glyeraldehyde 3-phosphate dehydrogenase (GAPDH) served as reference for normalization.

**Table 3.**
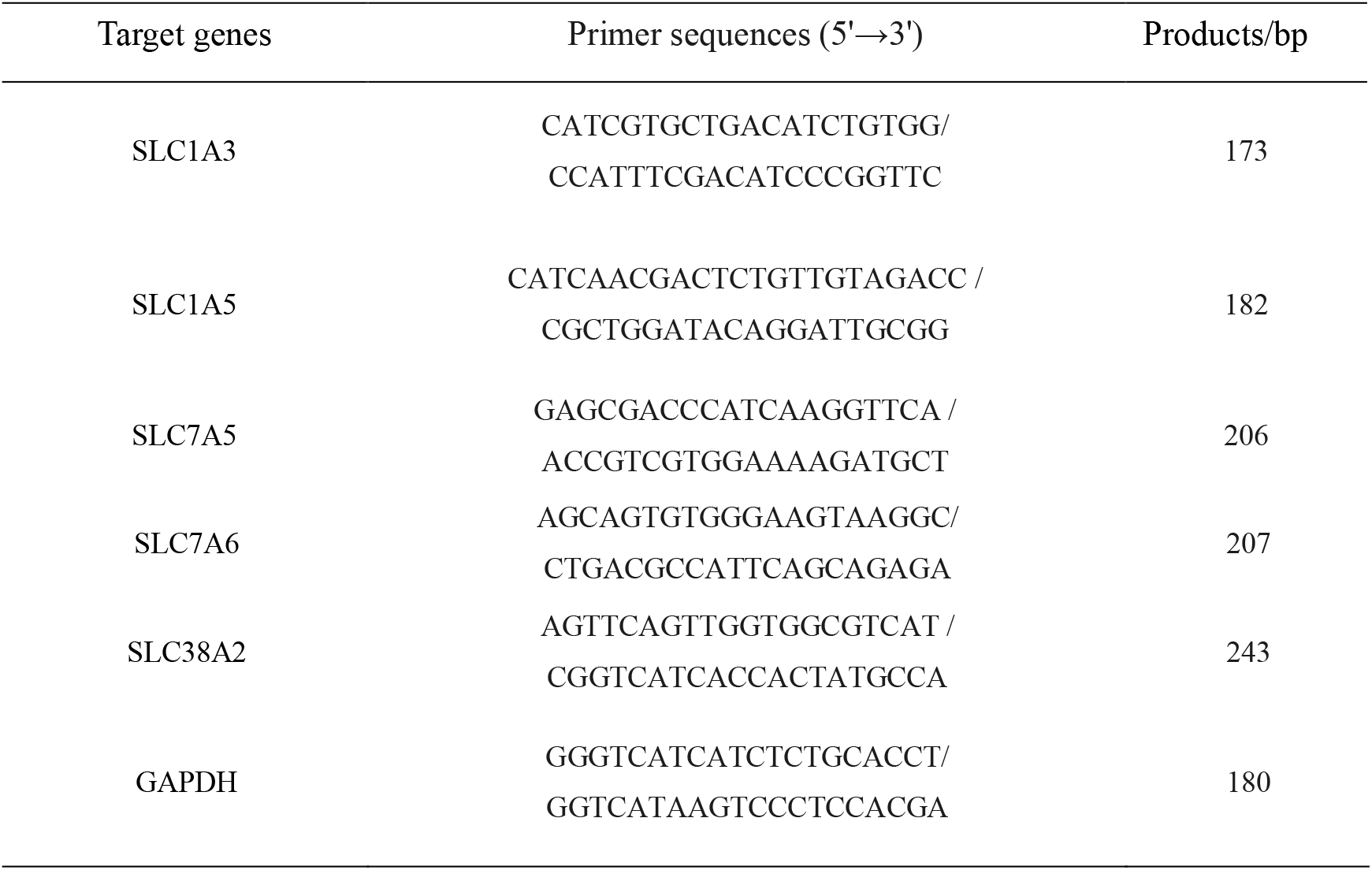
Primer sequences used for qRT-PCR analysis of target genes in lactating goats.

### Protein extraction from mammary tissues

Protein extraction from mammary tissues was performed according the method described by Duanmu et al (12). The mammary tissue samples of all goats in BG in equal quality were mixed and washed three times with ice-cold saline containing 1 mM PMSF, then homogenized in the ice-cold lysis buffer (2M thiourea, 7M urea, 50mM DTT, 2% (w/v) CHAPS, 0.5% (v/v) Bio-Lyte Ampholyte and 1mM PMSF) by 1:5 (w/v). The homogenates were kept at room temperature for 30 min, followed by centrifugation at 15,000 g for 30 min at 4 °C. The HG samples were treated in the same way. The samples were stored at −80 °C until analysis. The protein concentration of the supernatant was determined by RC DCTM (Bio-Rad, USA) kit.

### Two-dimensional gel electrophoresis (2-DE)

The operation method of 2-DE was referred to our laboratory previous research by Jiang et al (13). The first dimension used was isoelectric focusing (IEF). The extracted protein (1000 mg) was loaded onto the 17 cm IPG gel strips (nonlinear, pH 3.0-10.0, Bio-Rad, USA) according to Chen et al (14). Passive rehydration (13 h with 50 V). IEF was performed with a voltage gradient of 250 V for 1 h, 500 V for 1 h, 2000 V for 1 h, 8000 V for 3 h, followed by holding at 8000 V until a total of at least 60 000 V-h was reached. Then IPG strips were equilibrated by serial incubation for 15 min in equilibration buffer (6M urea, 30% (v/v) glycerol, 2% (w/v) SDS, 50 mM Tris-HCl (pH 8.8) and 1% (w/v) DTT) and in equilibration buffer containing 2.5% (w/v) iodoacetamide instead of 1% DTT. Equilibrated IPG strips were transferred onto the 12.5% SDS-PAGE for the second dimension (15). Gels were fixed in 12% trichloroacetic acid for 2 h, then stained with 0.08% (w/v) Coomassie Brilliant Blue G250 staining solution for 20 h. The excess of dye was removed with MilliQ water, and scanned with Molecular Imager (Versa Doc3000, Bio-Rad, USA). Standardization, background elimination, spot detection, gel matching and interclass analysis were performed as previously described using the PDQuest 8.0 software (Bio-Rad) (16). Three replicates were performed per sample. Protein spots were considered to be differentially expressed only if they showed 1.5-fold change in intensity, and satisfied the non-parametric Wilcoxon test (p< 0.05). Only the spots with the same changing trend in all three gels were considered for further analysis.

### Trypsin digestion and MS analysis

Selected gel spots were manually excised and washed twice with MilliQ water. Trypsin digestion test was performed as described by Ura et al (17). The digested proteins were air-dried and analyzed by using a 4800 MALDI-TOF/TOF proteomics analyzer (Applied Biosystems, USA). A protein spot digested with trypsin was used to calibrate the mass spectrometer. A mass range of 800-3500 Da was used. Combined search (MS plus MS/MS) was performed using GPS Explorer TM software v3.6 (Applied Biosystems, USA) and the MASCOT search engine (Matrix Science Ltd., UK). Proteins were considered as positive hints if at least two independent peptides were identified with medium (95%) or high (99%) confidence.

### Western blot analysis

Protein was extracted from mammary tissue samples using lysis buffer (Cell Signaling) plus PMSF (1 mM) and total protein was quantified by the bicinchoninic acid (BCA) assay (Pierce, Rockford, IL, USA). The protein of all samples in BG in equal quality were mixed, the HG samples were treated in the same way. Four replicates were performed per sample. We isolated 30 μg of protein from each sample, which was subjected to electrophoresis on SDS-PAGE. The separated proteins were transferred onto nitrocellulose membranes (Bio Trace, Pall Co., USA). The blots were incubated with the following Cell Signaling Technology primary antibodies for overnight at 4°C with a dilution of 1:1000 in block: rb-anti-Mammalian target of rapamycin (rb-anti-mTOR, #2983S), rb-anti-phospho-mTOR (rb-anti-p-mTOR, #5536S), rb-anti-P70 ribosomal protein S6 kinase (rb-anti-P70S6K, #9202S), rb-anti-phospho-P70S6K (rb-anti-p-P70S6K, #5536S), rb-anti-eukaryotic translation initiation factor 4E (rb-anti-eIF4E, #9742S), rb-anti-phospho-eIF4E (rb-anti-p-eIF4E, #9741S), rb-anti-eukaryotic elongation factor-2 kinase (rb-anti-eEF2K, #3692S), rb-anti-phospho-eukaryotic elongation factor-2 kinase (rb-anti-p-eEF2K, #3691S), rb-anti-eukaryotic elongation factor-2 kinase (rb-anti-eEF2, #2332S). The blot was incubated with ABclonal Technology primary antibody for overnight at 4°C with a dilution of 1:1000 in block: rb-anti-beta-casein protein (rb-anti-CSN2, #A12749S). The blot was incubated with primary antibody for overnight at 4°C with a dilution of 1:400 in block: rb-anti-lactoferrin (rb-anti-Lf, our laboratory) (18). A rb-anti-GAPDH primary antibody (a531, Bioworld, China, 1: 10,000) was also incubated with the blots to provide a reference for normalization. After washing the membranes, an incubation with HRP-conjugated secondary antibody was performed for 2 h at room temperature. Finally, the blots were washed and signal was detected by enhanced chemiluminescence (ECL) using the LumiGlo substrate (Super Signal West Pico Trial Kit, Pierce, USA). ECL signal was recorded using an imaging system (Bio-Rad, USA) and analyzed with Quantity One software (Bio-Rad, USA). The phosphorylation level of mTOR, P70S6K, eIF4E and eEF2K was determined by the ratio of p-mTOR to total mTOR, p-P70S6K to total P70S6K, p-eIF4E to total eIF4E and p-eEF2K to total eEF2K, respectively. The expression level of eEF2, CSN2 and LF were determined by the ratio of eEF2 to GAPDH, the ratio of CSN2 to GAPDH and the ratio of LF to GAPDH.

### Statistical analyses

Data were analyzed using the Statistical Package for Social Science (SPSS Inc., Chicago, IL, USA) and all data are presented as the mean ± SEM. The 2^−ΔΔCt^ method was applied to analyze the real-time PCR. Data were considered statistically significant if *P* < 0.05. The numbers of replicates used for statistics are noted in the Tables and Figures.

## Results

### Chromatographic separation of amino acid standard solution

A chromatogram of synthetic mixture of amino acid standards was received by RP-HPLC. Each peak represents one of specific amino acid, the peak was compact and symmetrical. 14 amino acids were completely separated (T<50 min) under the experimental conditions used, the peak sequence of 14 amino acids was: aspartic acid, glutamic acid, asparagine, serine, glutamine, glycine, arginine, alanine, tyrosine, valine, tryptophan, phenylalanine, isoleucine, leucine. We prepared the amino acid mixed standard solution at concentration of 31.25 - 500 μmol/L. After derivatization, we measured the samples. Taking the amino solution concentration (*x*) as the abscissa and corresponding peak area (*y*) as the coordinate. As shown in Table 4, in 31.25 - 500 μmol/L concentration range, amino acid standard concentration was linear related to the peak area and the correlation coefficients is 0.9960 - 0.9999. The Intra-day RSD and Inter-day RSD is between 2.15% - 4.05% and 3.13% - 4.83%, respectively, which are within 6%. These parameters results indicated that this sensitive procedure could be used for the quantitative analysis of amino acid in tissues.

**Table 4.** Determination parameters of standard amino acids.

| Amino acids | Retention time (t/min) | Regression equation (μmol/L) | Correlation coefficients |  | Intra-day | Inter-day |
| --- | --- | --- | --- | --- | --- | --- |
|  |  |  | Intra-day (r) |  | RSD (%) | RSD (%) |
| Asp | 6.025 | y = 1.0518x - 7.5833 | r=0.9998 |  | 2.52 | 3.14 |
| Glu | 8.624 | y = 1.1656x - 13.771 | r=0.9996 |  | 3.26 | 4.01 |
| Asn | 16.108 | y = 0.5021x + 4.6208 | r=0.9992 |  | 2.15 | 3.17 |
| Ser | 18.58 | y = 0.8926x - 6.2708 | r=0.9992 |  | 3.23 | 3.92 |
| Gln | 20.227 | y = 0.8177x + 1.5174 | r=0.9992 |  | 3.06 | 3.98 |
| Gly | 23.661 | y=1.1565x- 11.778 | r=0.9987 |  | 2.88 | 3.16 |
| Arg | 25.031 | y=1.271x- 21.429 | r=0.9974 |  | 2.69 | 3.64 |
| Ala | 30.285 | y = 0.9363x - 8.8435 | r=0.9988 |  | 2.67 | 3.13 |
| Tyr | 32.45 | y=1.546x- 10.704 | r=0.9985 |  | 3.59 | 3.78 |
| Val | 39.702 | y = 1.7831x - 2.5696 | r=0.9990 |  | 3.23 | 4.23 |
| Trp | 40.734 | y = 1.3565x - 48.778 | r=0.9979 |  | 3.18 | 3.67 |
| Phe | 42.086 | y = 1.6486x - 45.243 | r=0.9975 |  | 3.89 | 4.35 |
| Ile | 43.492 | y = 1.7844x - 21.226 | r=0.9977 |  | 4.01 | 4.54 |
| Leu | 44.521 | y = 1.0609x + 7.7087 | r=0.9986 |  | 4.05 | 4.83 |

Note: Asp, aspartic acid; Glu, glutamic acid; Asn, asparagines; Ser, serine; Gln, glutamine; Gly, glycine; Arg, arginine; Ala, alanine; Tyr, tyrosine; Val, valine; Trp, tryptophan; Phe, phenylalanine; Ile, isoleucine; Leu, leucine. RSD, relative standard deviation. The peak times of Met and Val, Lys and Tyr are almost the same, so Val and Tyr are selected in this experiment, Met and Lys are removed. The peak times of Thr and Arg are very close, and compared with Arg, the peak of Thr is very low and can not be completely separated, so Arg is selected. Cys peak is very low and the quantitative error is large, so it is also removed. Under this condition, His has two peaks, the exact peak time can not be determined, so the His is removed. Pro does not contain primary ammonia, and the derivatizing agent phthalaldehyde cannot be derivatized, so it cannot be determined.

### Amino acid content in jugular venous plasma

As shown in Figure 1, all of the 14 kinds of amino acid concentrations were higher in buffering agent groups than that in the high grain group. The amino acid content of Gln showed very significant difference (*P* < 0.01) between these two groups. And the amino acid concentrations of Asn, Ser, Gly and Arg were significantly higher (*P* < 0.05) than that in high grain group.

**Figure 1.**
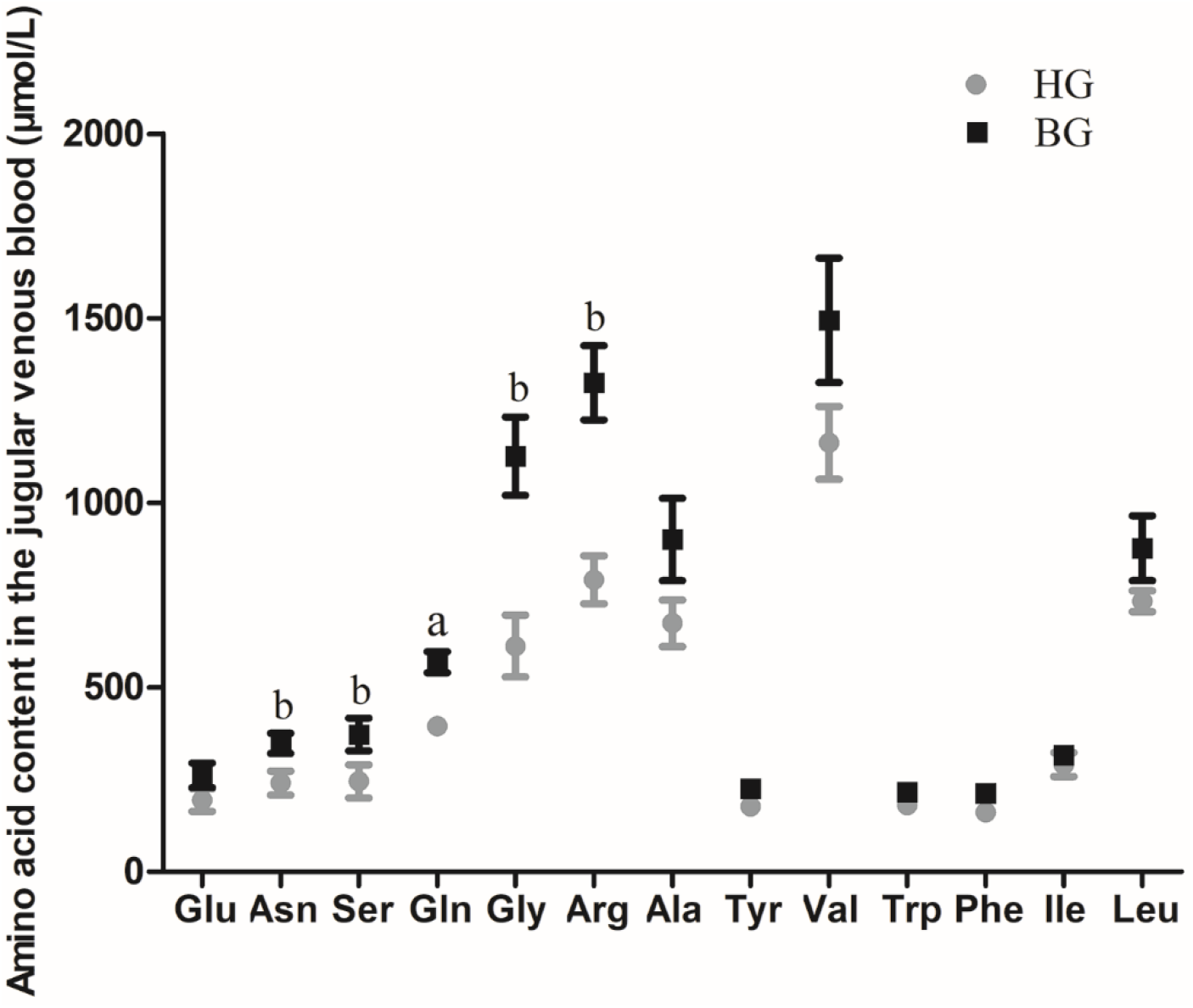
Impact of high-grain diet with buffering agent on amino acid content in plasma of lactating goats. Values are mean ± SEM, n = 6/group. ^b^p<0.05 and ^a^p<0.01, compared with high grain group.

### Different types of amino acids in jugular venous plasma

As shown in Figure 2, the amino acid concentrations of TFAA, GAA and NEAA were significantly higher (*P* <0.05) than that in high grain group (2A). And the total AA content performed by Total Amino Acid assay kit in plasma of lactating goats showed very significant difference (*P* <0.01) between these two groups (2B).

**Figure 2.**
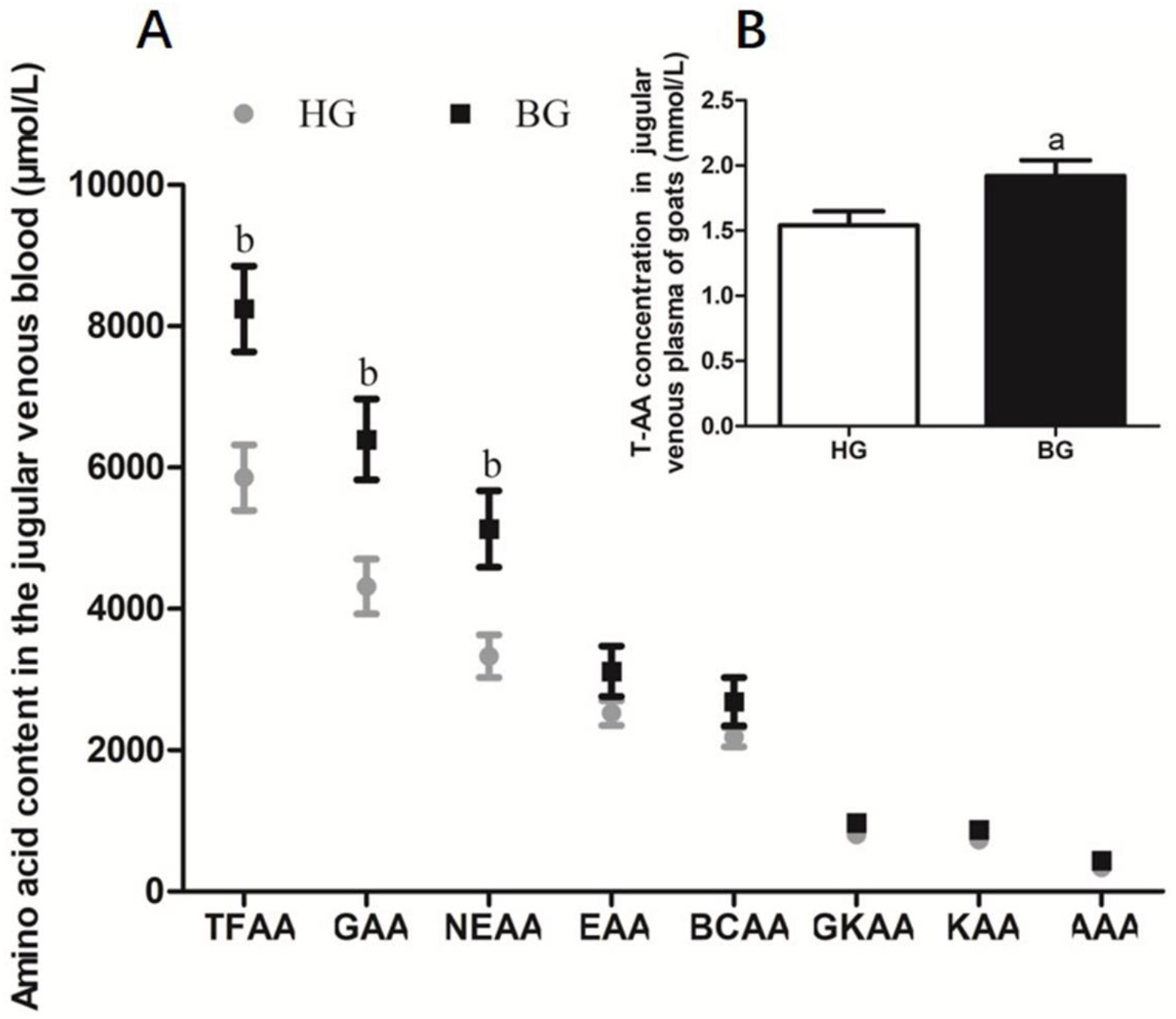
Different types of amino acids content in plasma of lactating goats. (A) Impact of high-grain diet with buffering agent on different types of amino acids content in plasma of lactating goats by HPLC. (B) Impact of high-grain diet with buffering agent on total amino acids content in plasma of lactating goats by Total Amino Acid assay kit. Values are mean ± SEM, n = 6/group. ^b^p<0.05 and ^a^p<0.01, compared with high grain group. TFAA, total free amino acid; GAA, glycogenic amino acid; NEAA, non-essential amino acid; EAA, essential amino acid; BCAA, branched-chain amino acid; GKAA, glucogenic and ketogenic amino acid; KAA, ketogenic amino acid; AAA, aromatic amino acid.

### mRNA transcription level of amino acid transport

The excitatory amino acid transporter 1 (EAAT1, encoded by gene SLC1A3), alanine-serine-cysteine transporter 2 (ASCT2, encoded by gene SLC1A5), L-type amino acid transporter 1 (LAT1, encoded by gene SLC7A5), sodium-independent neutral and basic amino acid transporter (y^+^LAT2, encoded by gene SLC7A6) and sodium-coupled neutral amino acid transporter 2 (SNAT2, encoded by gene SLC38A2) were reported as transporters of amino acids to maintain cell growth and protein synthesis in ruminant (19–21). However, the function of these amino acid transporters in regulating milk protein synthesis in the mammary gland of the lactating goats remains largely unknown. Therefore, mRNA expression levels of the corresponding genes encoding amino acid transporters in the mammary gland were detected by RT-PCR to reflect the role of these amino acid transporters in the mammary gland. SLC1A3, SLC1A5, SLC7A5, SLC7A6 and SLC38A2 mRNA expression levels in mammary tissues were up-regulated in BG goats compared to HG goats, the mRNA expressions of SLC38A2 was significantly increased (*P* < 0.05), and the mRNA expressions of SLC7A6 was very significantly increased (*P* < 0.01) in BG goats compared to HG goats (Figure 3).

**Figure 3.**
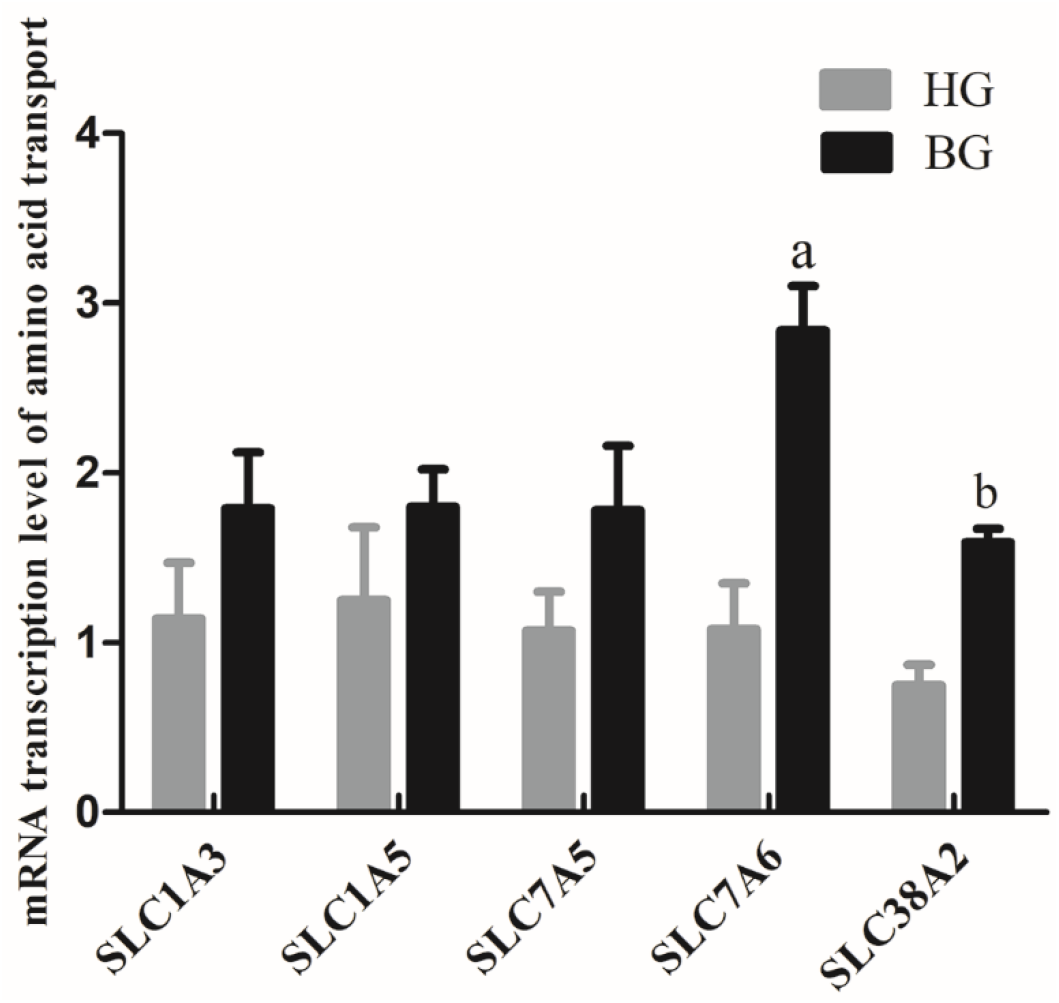
mRNA expressions of amino acid transport SLC1A3, SLC1A5, SLC7A5, SLC7A6 and SLC38A2. Each sample was first normalized against its GAPDH transcript level, and then normalized to the HG. In order to calculate differences in the expression level of each target gene, the 2^−ΔΔCt^ method for relative quantification was used, according to the manufacturer’s manual. Values are mean ± SEM. n = 6/group. ^b^p<0.05 and ^a^p<0.01, compared with high grain group.

### Global identification of differentially expressed proteins in the mammary gland

Comparative proteomic analysis was performed between HG and BG lactating Saanen goats’ mammary tissues, in order to understand the influence of adding buffering agent in high-grain diet on the mammary metabolism. As shown in Figure 4, an average of 1200 spots were detected on gels for both types of proteomes. We successfully found a total of 55 differential protein spots and 32 differential protein spots (p < 0.05; in terms of expression, all 32 with a fold change ≥1.5-fold) were successfully identificated in BG vs. HG using 2-DE technique and MALDI-TOF/TOF proteomics analyzer. Of these, 15 proteins showed increased expression and 17 proteins showed decreased expression in BG vs. HG respectively.

**Figure 4.**
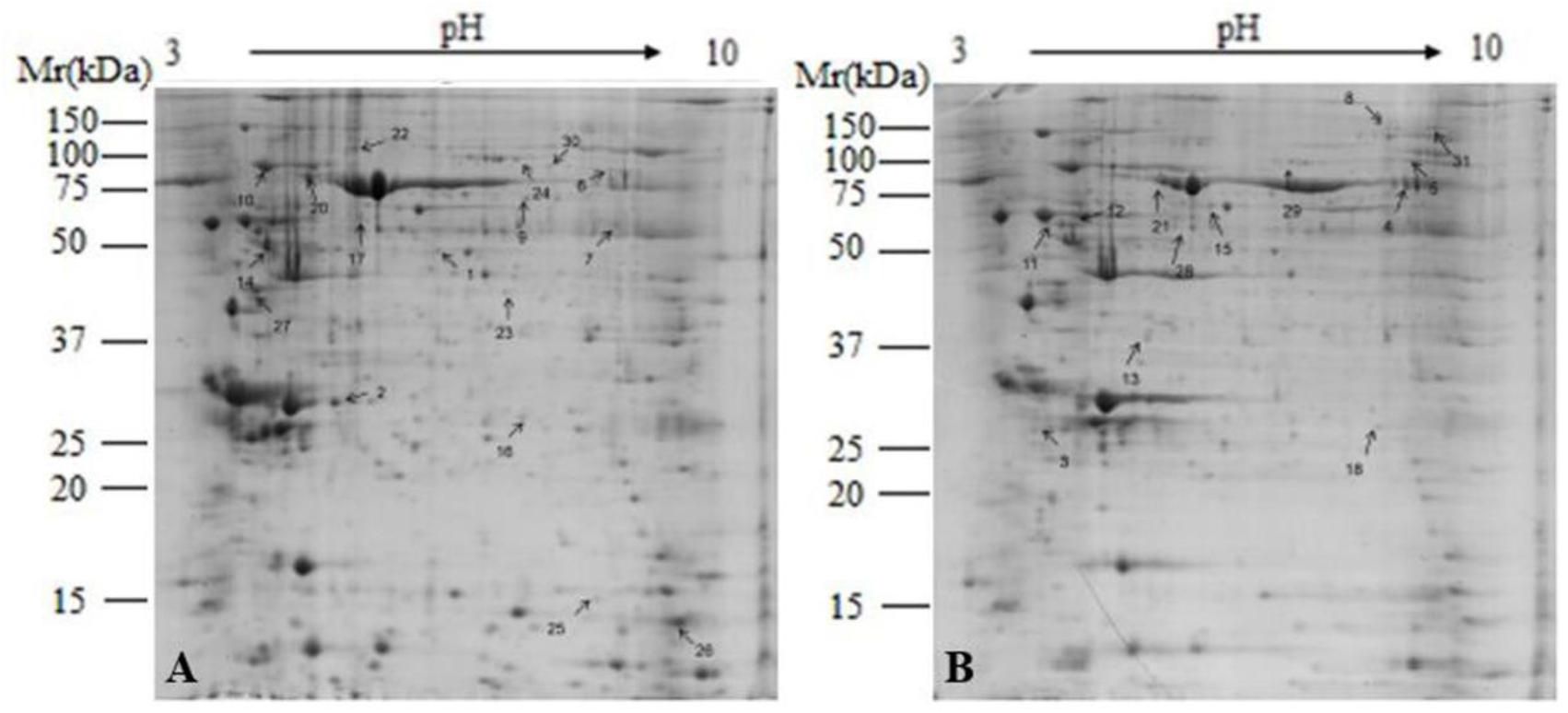
Representative 2-DE images of proteins extracted from lactating goat mammary gland. (A) High-grain group; (B) Buffering agent group. Equal amounts of protein (1000 mg) were loaded onto the 17 cm IPG gel strip (nonlinear, pH 3.0-10.0), and separated on 17-cm IPG strips (pH 3.0-10.0), followed by electrophoresis on 12.5% SDS-PAGE gels for second dimension electrophoresis. Black arrows indicate differential protein spots (≥ 1.5-fold).

We can observe the main differential protein spots related to amino acid metabolism, glucose metabolism, lipid metabolism, oxidative stress, mitochondrial function, cytoskeletal structure and immune protein depicted in Table 5.

**Table 5.** Differential expression protein spots by MAIDI-TOF-TOF.

| Spot no. | Identified protein name | Gene symbol | Accession no. | Mr(kDa) | pI | Protein expression | 1.5Fold change |
| --- | --- | --- | --- | --- | --- | --- | --- |
| Amino acid metabolism |  |  |  |  |  |  |  |
| 1 | Ornithine aminotransferase | OAT | gi\|803207295 | 37.8 | 8.37 | Down | 0.65 |
| 2 | Beta casein | CSN2 | gi\|1211 | 24.9 | 5.26 | Up | 2.73 |
| 3 | Peptidylprolyl isomerase A | PPIA | gi\|410066839 | 18.0 | 8.34 | Down | 0.66 |
| 4 | Lactoferrin | LF | gi\|56544486 | 79.2 | 8.4 | Up | 2.27 |
| Glucose Metabolism |  |  |  |  |  |  |  |
| 5 | Pyruvate kinase PKM isoform X7 | PKM | gi\|426232638 | 58.5 | 7.60 | Down | 0.23 |
| 6 | UTP--glucose-1-phosphate uridylyltransferase isoform X1 | UGP2 | gi\|426223460 | 57.0 | 7.68 | Up | 2.13 |
| 7 | Transketolase | TKT | gi\|426249391 | 68.5 | 7.2 | Up | 2.52 |
| 8 | Aconitate hydratase, mitochondrial isoform X1 | ACO2 | gi\|803058827 | 95.9 | 6.78 | Down | 0.5 |
| 9 | Aldehyde dehydrogenase, mitochondrial | ALDH2 | gi\|426247368 | 57.1 | 7.55 | Down | 0.52 |
| 10 | Glucose-regulated protein | HSPA5 | gi\|426223038 | 72.5 | 5.07 | Up | 1.66 |
| Cytoskeletal structure |  |  |  |  |  |  |  |
| 11 | Tubulin | Tubulin | gi\|257097261 | 50.2 | 4.78 | Up | 1.66 |
| 12 | Vimentin | VIM | gi\|145226795 | 53.7 | 5.02 | Up | 1.56 |
| 13 | F-actin-capping protein subunit beta isoform X1 | CAPZB | gi\|426222052 | 34.0 | 5.82 | Down | 0.49 |
| 14 | Protein disulfide-isomerase A6 | PDIA6 | gi\|803229999 | 48.5 | 4.95 | Up | 2.1 |
| Lipid metabolism |  |  |  |  |  |  |  |
| 15 | Glycerol kinase isoform X5 | GK | gi\|426256814 | 61.4 | 5.60 | Down | 0.41 |
| 16 | Acid synthase isoform X2 | FASN | gi\|803208475 | 27.6 | 5.93 | Up | 6 |
| Oxidative stress |  |  |  |  |  |  |  |
| 17 | Cytochrome b-c1 complex subunit 1 | UQCRC1 | gi\|803186314 | 51.9 | 6.03 | Down | 0.42 |
| 18 | Glutathione S-transferase P | GSTP1 | gi\|803225108 | 23.8 | 6.89 | Up | 1.96 |
| Mitochondrial function |  |  |  |  |  |  |  |
| 19 | ATP synthase subunit beta, mitochondrial | ATP5B | gi\|426224929 | 56.2 | 5.15 | Up | 1.66 |
| 20 | NADH-ubiquinone oxidoreductase 75 kDa subunit | NDUFS1 | gi\|426221412 | 80.4 | 5.90 | Down | 0.46 |
| Immune protein |  |  |  |  |  |  |  |
| 21 | Pre-pro serum albumin | ALB | gi\|1387 | 71.1 | 5.80 | Down | 0.18 |
| 22 | Polymeric immunoglobulin receptor | PIGR | gi\|426239425 | 83.7 | 5.79 | Down | 0.17 |
| 23 | Annexin A1 | ANXA1 | gi\|426220300 | 39.0 | 6.17 | Down | 0.47 |
| 24 | Serotransferrin isoform X2 | TF | gi\|803335974 | 79.4 | 6.31 | Up | 2.38 |
| 25 | Beta globin chain | HBB | gi\|86129745 | 16.0 | 6.75 | Up | 1.89 |
| 26 | Alpha globin chain | HBAI | gi\|1787 | 15.3 | 8.72 | Up | 1.60 |
| Other |  |  |  |  |  |  |  |
| 27 | Protein FAM186B isoform X1 | FAM186B | gi\|803043984 | 111.3 | 8.87 | Down | 0.47 |
| 28 | T-complex protein 1 subunit epsilon | CCT5 | gi\|426246714 | 60.03 | 5.6 | Down | 0.57 |
| 29 | Radixin isoform X1 | RDX | gi\|803154243 | 68.4 | 6.03 | Down | 0.53 |
| 30 | Lamin isoform X2 | LMNA | gi\|803255736 | 74.4 | 6.76 | Down | 0.48 |
| 31 | Elongation factor 2 isoform X2 | EEF2 | gi\|426229147 | 96.2 | 6.41 | Down | 0.36 |
| 32 | cytochrome b-c1 complex subunit 2 | UQCRC2 | gi\|426254425 | 48.8 | 8.89 | Up | 2.47 |

Beta casein (CSN2) is the main composition of milk protein. Lactoferrin (LF) is a nonheme iron binding glycoprotein in milk and a member of transferring family, during lactation expressed and secreted by the mammary epithelial cells at mucosal surface.

The protein expression of CSN2 and LF were up-regulated by 2-DE technique and MALDI-TOF/TOF proteomics analyser in BG as compared to the HG.

### Validation of differentially expressed milk proteins

To further validation the CSN2 and LF proteins (main composition of milk protein), we performed western blot analysis for verification (Figure 5). The CSN2 protein expression level was extremely significant higher (*P* < 0.01) in BG than that in HG. And the LF protein expression level was also higher in BG than that in HG (*P* < 0.05).

**Figure 5.**
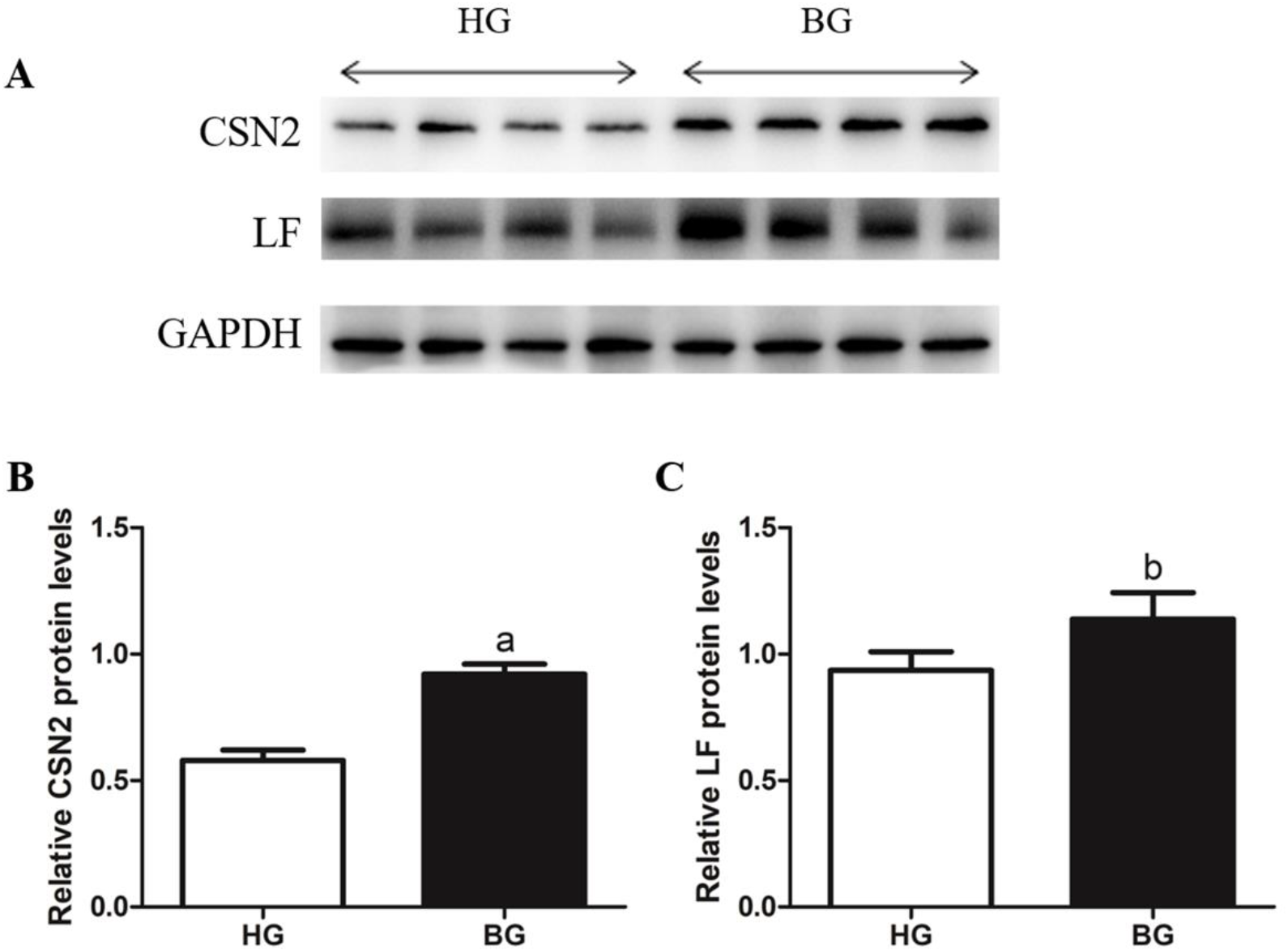
Western blot analysis of the CSN2 and LF. (A) Indicated protein levels were measured by western blotting analysis. (B and C) Relative protein levels of CSN2 (B) and LF (C) from the western blots were quantified by gray scale scan. Values are mean ± SEM. n = 4/group. ^b^p<0.05 and ^a^p<0.01, compared with high grain group.

### The signaling pathway for milk protein synthesis

Western blot was applied to study the mechanism of the milk protein synthesis, the mTOR phosphorylation ratio and P70S6K phosphorylation ratio were significant up-regulated (*P* < 0.05, Figure 6B and 6C), the eIF4E phosphorylation ratio and the expression level of eEF2 were up-regulated, and showed significant difference (*P* < 0.01, Figure 6D and 6F), but the eEF2K phosphorylation ratio was down-regulated (*P* < 0.01, Figure 6E). It suggested that adding buffering agent could promote the protein expression levels of mTOR phosphorylation ratio, P70S6K phosphorylation ratio, eIF4E phosphorylation ratio and eEF2, and suppress the protein expression of eEF2K phosphorylation ratio.

**Figure 6.**
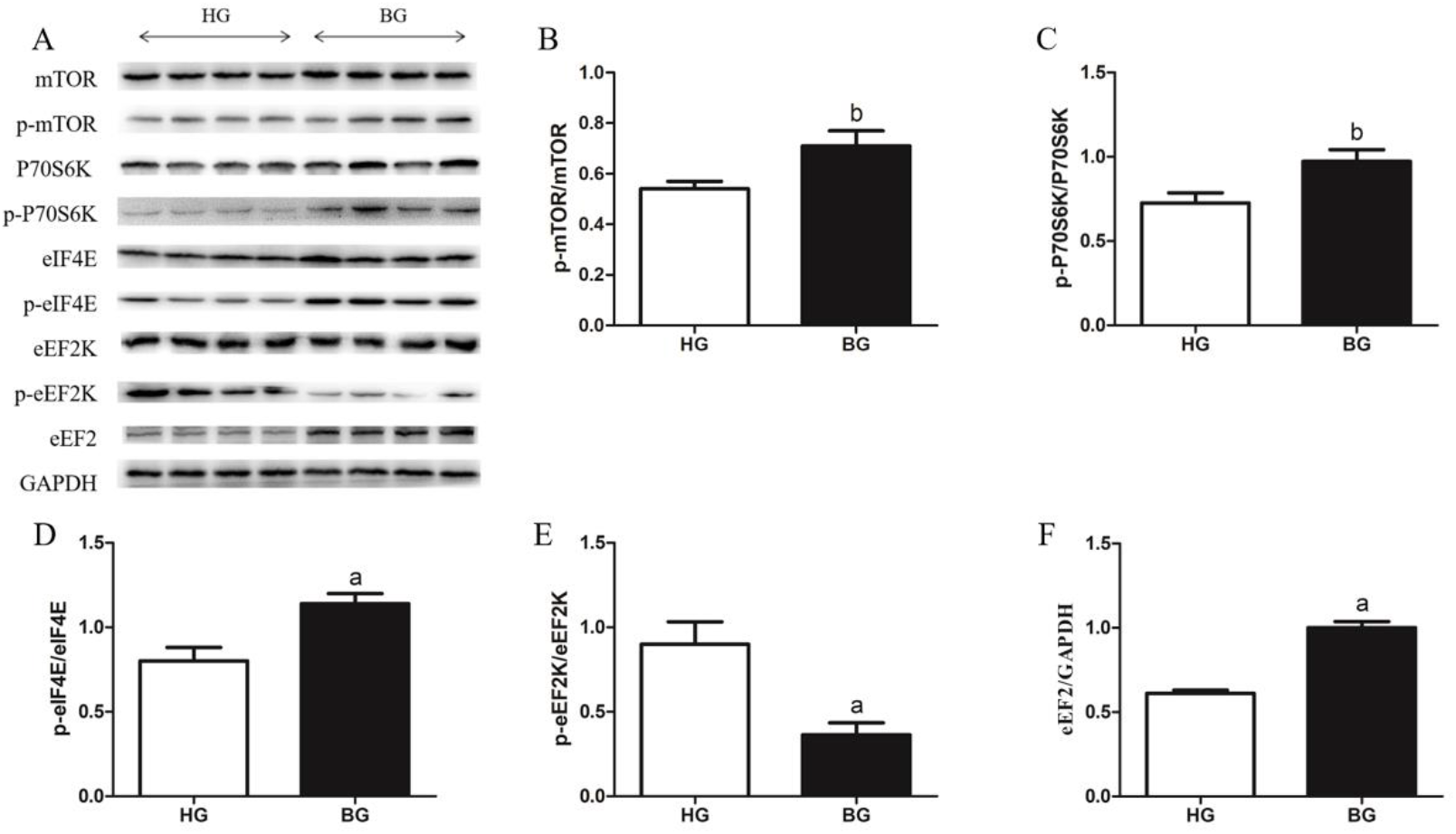
Milk protein synthesis mTOR related signaling pathway protein expressions. (A) mTOR related signaling pathway protein expression levels were measured by western blotting analysis. (B) The ratios of phosphorylated to total mTOR were quantified by gray scale scan. (C) The ratios of phosphorylated P70S6K to total P70S6K were quantified by gray scale scan. (D) The ratios of phosphorylated eIF4E to total eIF4E were quantified by gray scale scan. (E) The ratios of phosphorylated eEF2K to total eEF2K were quantified by gray scale scan. (F) Relative protein levels of eEF2 from the western blots were quantified by gray scale scan. Values are mean ± SEM. n = 4/group. ^b^p<0.05 and ^a^p<0.01, compared with high grain group.

## Discussion

Milk production of dairy is determined mainly by the fat synthesis and milk protein and proliferation abilities of mammary epithelial cells (BMECs) (22). And milk proteins in the lactating ruminant mammary gland are primarily synthesized from circulating plasma amino acids, amino acids present in both free and peptide-bound forms, used by tissues both as building blocks for protein synthesis and as signaling molecules to regulate the protein synthetic machinery (23,24). The mammary gland requires large amounts of amino acids for the synthesis of milk protein. Uptake of amino acids from the arterial blood of the lactating dam is the ultimate source of proteins (primarily β-casein and α-lactalbumin) in milk (25). During lactation the ruminant mammary gland must import from the plasma and endogenously synthesize sufficient quantities of amino acids to elevate milk protein synthesis (26). Identifying and understanding changes of amino acids in plasma and and amino acids transport into the lactating mammary gland may provide fundamental knowledge towards the development of nutritional regimes aimed at elevating milk production. Here we found that compared with the HG, 14 kinds of free amino acid concentrations determined were higher in BG, and the amino acid concentrations of TFAA, GAA and NEAA were significantly higher (*P* <0.05) than that in high grain group by HPLC. And the total AA concentration performed by Total Amino Acid assay kit in plasma of lactating goats also showed very significant difference (*P* <0.01) between these two groups. Therefore, adding buffering agent could promote the increase of amino acid concentration in blood compared with long-term feeding of high grain diet.

Presently, the amino acid transporter system in the mammary gland is not well-understood, but information suggests that the mammary gland has a transport system similar to other organs (intestine, kidney, placenta) (27). The future studies need to be conducted to identify changes in mammary genes involved in amino acid transport and uptake in the mammary gland. Main transporters involved in amino acid transport and uptake in lactating goat’s mammary gland were investigated. SLC1A3, SLC1A5, SLC7A5, SLC7A6 and SLC38A2 mRNA expression levels in the mammary gland were detected by RT-PCR to reflect the role of these amino acid transporters in the mammary gland. We found that the mRNA expression levels of SLC1A3, SLC1A5, SLC7A5, SLC7A6 and SLC38A2 in mammary tissues were up-regulated in BG goats compared to HG goats, the mRNA expressions of SLC38A2 was significantly increased (*P* < 0.05), and the mRNA expressions of SLC7A6 was very significantly increased (*P* < 0.01) in BG goats compared to HG goats. EAAT1, encoded by SLC1A3 is one isoform of system X_AG_^−^ and is a Na^+^-dependent transporter with high affinity for Asp and Glu. ASCT2, encoded by SLC1A5 is Na^+^-dependent, and has affinity for small neutral AA, such as Ala, Ser and Cys (28). y^+^LAT2, encoded by SLC7A6 is a catalytic light chain (y^+^LAT) and togethers with a heavy subunit (4F2hc) linked by a disulfide bond to form a heteromeric Na^+^-dependent transporters y^+^LAT2/4F2hc (Torrents et al. 1998) (29). y^+^LAT2/4F2hc is located in the basolateral cell membrane, functions as obligatory asymmetric AA exchangers and has affinity for Lys, Arg, Gln, His, Met, Leu, Ala and Cys (30). LAT1, encoded by SLC7A5 found in many different types of mammalian cells, is indispensable as a transporter of essential AA to maintain cell growth and protein synthesis (31). SNAT2, encoded by SLC38A2 is expressed in the mammary gland and plays an important role in the uptake of alanine and glutamine which are the most abundant amino acids transported into this tissue during lactation, and has affinity for Gly, Pro, Ala, Ser, Cys, Gln, Met, His and Asn. The expression of SNAT2 can be upregulated by amino acids and hormones (prolactin and 17β-estradiol) (32). Our experimental data, combined with these reports, suggest that adding buffering agent could not only promote the increase of amino acid concentration in blood but also through amino acid transporters transport more amino acids from the blood to the mammary gland for the synthesis of milk protein compared with long-term feeding of high grain diet.

Milk protein is mainly composed of casein and whey protein, the protein content in milk from caprine species is approximately 2.1-3.5%, caprine milk has a high casein to whey ratio of 6.0, casein is synthesized by the epithelial cells of the mammary gland (33). It is a group of phosphoproteins unique to milk and contains a large amount of phosphorus and calcium. The casein fraction consists of α_s1_-, α_s2_-, β- and κ-casein. The whey fraction consists of mainly β-lactoglobulin (β-lg), α-lactalbumin (α-la), immunoglobulins, lactoferrin (LF) and lysozyme (34–36). In order to verify the amino acids go into the mammary gland are more utilized by the mammary gland to synthesize milk proteins, we used 2-DE technique and MALDI-TOF/TOF proteomics analyzer to confirm it. We observed the protein expressions of CSN2 and LF were up-regulated by 2-DE technique and MALDI-TOF/TOF proteomics analyser in BG as compared to the HG. Further verification by WB indicated that CSN2 protein expression level was extremely significant higher (*P* < 0.01) in BG than that in HG. And the LF protein expression level was also higher in BG than that in HG (*P* < 0.05).

It has been reported that β-casein has the highest content in casein of caprine milk, which is different from the milk of bovine (37). Lactoferrin is a nonheme iron binding glycoprotein in milk and a member of transferring family, during lactation expressed and secreted by the mammary epithelial cells at mucosal surface. The results showed that adding buffering agent in high grain diet promoted the synthesis of casein which is the main components of milk protein and lactoferrin which comes from whey protein in the mammary gland, so it was further verified that more milk protein, especially casein and lactoferrin was synthesized by amino acid in blood entering into the mammary gland.

It is widely accepted that mammalian target of rapamycin (mTOR) is variety of components of protein synthesis and a key regulator of milk protein synthesis, most reports were concerned with the role of amino acids and mTOR in protein synthesis, but the mechanism of how amino acids enter the mammary gland to regulate the mTOR signaling pathway and promote milk protein synthesis is still not very clear (38–40). To further explore the mechanisms by which the buffering agent treatment regulated related proteins expression in milk protein synthesis, we studied the activity of the mTOR signaling pathway. In the present study, the results showed that the amino acids could activate mTOR signaling which is consistent with previous reports. Here we identifies mTOR phosphorylation site as an indicator of activated mTOR pathway, which was increased by the increasing amino acids entering into mammary gland. mTORC1 activates P70S6K via phosphorylation, which in turn activates several proteins that contribute to increases in protein synthesis. In addition, activation of P70S6K inhibits eEF2K via phosphorylation, subsequently stimulating eEF2, which activates translocation elongation. When mTORC1 is phosphorylated it inhibits 4E-binding protein 1 (4EBP1) through phosphorylation, which inhibits eIF4E. eIF4E activates cap-dependent translocation increasing protein synthesis. Through these various proteins, mTORC1 stimulates cell growth by increasing cap-dependent translocation, translation elongation, mRNA biogenesis, and ribosome biogenesis, which leads to an increase in overall protein synthesis.

## Conclusion

These results suggested that feeding of high-grain diet with buffering agent promoted the jugular vein blood of amino acids concentration, and the mRNA expressions of amino acid transport SLC1A3, SLC1A5, SLC7A5, SLC7A6 and SLC38A2 in mammary tissues were up-regulated, it showed that more amino acids flowed into the mammary. Two-dimensional electrophoresis and WB analysis further verification showed milk protein synthesis was increased. The results of the WB analysis suggested that the increase of milk protein synthesis was related to the activation of mTOR pathway signaling (Figure 7).

**Figure 7.**
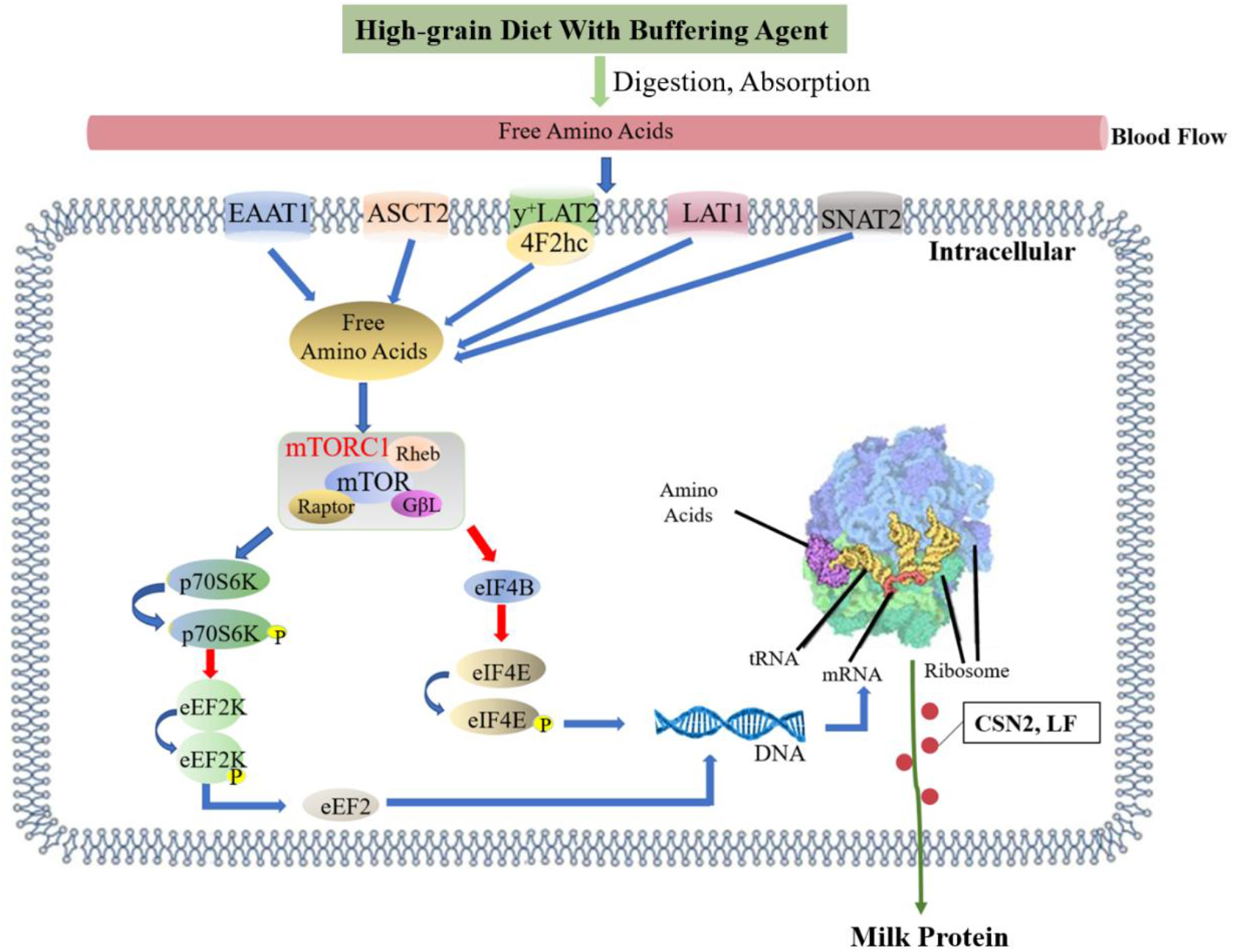
Effects of high-grain diet with buffering agent on the milk protein synthesis. Feeding of high-grain diet with buffering agent promoted the jugular vein blood of amino acids concentration, and the mRNA expressions of amino acid transport SLC1A3, SLC1A5, SLC7A5, SLC7A6 and SLC38A2 in mammary tissues were up-regulated, it showed that more amino acids flowed into the mammary. Amino acids in mammary gland activated mTOR pathway. mTORC1 activates P70S6K via phosphorylation, which in turn activated several proteins that contributed to promote protein synthesis. In addition, activation of P70S6K inhibited eEF2K via phosphorylation, subsequently stimulated eEF2, which activated translocation elongation. When mTORC1 was phosphorylated, it inhibited 4EBP1 through phosphorylation, which inhibits eIF4E. eIF4E activates cap-dependent translocation increasing protein synthesis. The blue arrow represents promotion, and the red arrow represents inhibition.

## Abbreviations

HPLC: high performance liquid chromatography
SARA: subacute ruminal acidosis
CSN2: beta casein
LF: Lactoferrin

## Acknowledgments

We are grateful to Hongyu Ma in Nanjing Agricultural University for advice on MS analysis. We also thank Xiangzhen Shen, Yingdong Ni and Su Zhuang (Nanjing Agricultural University) for their suggestions about the diet treatment.

## Authors’ contributions

MH conceived of the study, carried out the experiments and drafted the manuscript. XN and HW collected the sample and performed the research, analyzed data. MH and XN assisted with the sample analysis. YZ participated in the study’s design and coordination. All authors read and approved the final manuscript.

## Availability of data and materials

All data generated or analyzed during this study are included in this published article.

## Ethics approval and consent to participate

All animal procedures were approved by the Institutional Animal Care and Use Committee of Nanjing Agricultural University. The protocols were reviewed and approved, and the project number 2011CB100802 was assigned. The slaughter and sampling procedures strictly followed the ‘*Guidelines on Ethical Treatment of Experimental Animals*’ (2006) no. 398 established by the Ministry of Science and Technology, China and the ‘*Regulation regarding the Management and Treatment of Experimental Animals*’ (2008) no. 45 set by the Jiangsu Provincial People’s Government.

## Consent for publication

Not applicable.

